# Tropomyosin 3.1 Regulates Chondrocyte Cortical F-actin Stability and Homeostasis

**DOI:** 10.64898/2026.10.23.761071

**Authors:** Marin Herrick, Mark Arranguez, Josalyn Stack, Justin Parreno

## Abstract

**Objective:** Osteoarthritis is a degenerative joint disease of which the mechanisms and proteins contributing to articular cartilage breakdown have not yet been fully elucidated. In osteoarthritis, chondrocyte F-actin loss coincides with altered chondrocyte homeostasis favoring catabolism. While the loss of F-actin has been implicated as a driver of osteoarthritis progression, the mechanisms by which F-actin is lost and regulates a catabolic shift in gene expression require elucidation. We hypothesized that destabilizing F-actin by targeting inhibition of specific F-actin stabilizer Tropomyosins (Tpms) dysregulates chondrocyte homeostasis through regulation of actin-associated transcription factors, Yes-associated protein (YAP) and Myocardin-related transcription factor A (MRTF).

**Methods:** Using RT-PCR and Sanger sequencing, we determined the expression of Tpm isoforms present in cartilage. We then focused on the regulation of chondrocyte cortical F-actin by the specific Tpm isoform, Tpm3.1. Using confocal imaging for F-actin, we examined the effect of Tpm3.1 knockout in mouse femoral head cartilage and pharmacological inhibition (TR100) of primary bovine chondrocytes. Next, we examined the nuclear translocation of actin-associated regulators, YAP and MRTF, by TPM3.1 inhibition in primary chondrocytes. Finally, we determined the effect of YAP and MRTF inhibition on the regulation of chondrogenic and degradative enzyme gene expression in primary chondrocytes.

**Results:** Our results revealed TPM3.1 is present and localized cortically in mouse and bovine chondrocytes. There was a reduction in cortical F-actin staining in Tpm3.1 knockout mouse femoral head cartilage as well as in primary bovine chondrocytes treated with TR100. In primary chondrocytes, TR100 caused a catabolic shift in mRNA levels. This coincided with a decrease in the nuclear-to-cytoplasmic ratio of both YAP and MRTF in TR100-treated cells. Furthermore, pharmacological inhibition of either YAP or MRTF partially recapitulated the catabolic shift seen in TR100-treated chondrocytes by a reduction of chondrogenic mRNA levels.

**Conclusion:** Collectively, these results provide insight into the regulation of chondrocyte homeostasis by cortical F-actin stability and the downstream mediators of transcription. A greater understanding of the actin-associated molecules that regulate chondrocyte homeostasis in osteoarthritis may lead to novel insights to target disease progression.

## Introduction

Osteoarthritis is a degenerative disease characterized by articular cartilage breakdown. Current treatments are primarily symptomatic, with limited options to prevent progression. Since the mechanisms and regulatory molecules driving cartilage breakdown have not been fully elucidated, investigating the contribution of regulatory pathways, such as actin-mediated chondrocyte homeostasis, may provide insights into slowing disease progression before total joint replacement.

A catabolic shift in chondrocyte homeostasis is a major contributor to degeneration in osteoarthritis[1]. Healthy chondrocytes maintain the extracellular matrix by producing molecules including type II collagen (COL2), aggrecan (ACAN), and proteoglycan-4 (PRG4). During osteoarthritis, chondrocytes acquire a catabolic phenotype, downregulating matrix production[2–4] and elevating degradative enzyme expression (i.e., matrix metalloproteinases; MMPs)[5, 6]. Although the cytoskeleton regulates chondrocyte phenotype[7], the mechanisms by which cytoskeletal reorganization drives this catabolic shift remain unclear.

The actin cytoskeleton is a regulator of chondrocyte phenotype and function[7–10], existing in two forms: monomeric globular (G-) actin or polymerized filamentous (F-) actin. Native chondrocytes arrange F-actin cortically along the cell membrane[11]. While F-actin stress fiber formation has been associated with the degenerative chondrocyte phenotype[9], our recent surgical (destabilization of the medial meniscus) animal model showed that cortical F-actin loss coincides with degeneration[12]. Supporting this, human osteoarthritic chondrocytes and chondrocytes treated with inflammatory cytokines exhibit reduced cortical F-actin[13–15]. Furthermore, *in vitro* F-actin depolymerization reduces *Prg4*, *Col2*, and *Acan* expression while increasing expression of *Mmp-2*, *-3*, and *-13*[8, 16–18]. These observations suggest cortical F-actin loss drives cartilage degeneration. However, the mechanisms maintaining cortical F-actin stability remain unclear. Identifying the proteins that stabilize cortical F-actin and determining how F-actin destabilization alters downstream signaling are important for understanding how actin regulates chondrocyte homeostasis.

Tropomyosins (TPMs) are a large family of F-actin stabilizing proteins that organize F-actin into functionally distinct networks by coordinating its interactions with other actin-binding proteins. Over 40 non-redundant TPM isoforms exist, each specifying distinct cellular F-actin organization[19]. We identified Tpm3.1 in human and bovine passaged chondrocytes[8], and found that while TPM3.1 regulates stress fiber stability in passaged chondrocytes, it associates with cortical F-actin in primary chondrocytes[8]. However, whether TPM3.1 maintains cortical F-actin remains unknown. Because TPM3.1 stabilizes cortical F-actin in other cell types[20, 21], is mechanosensitive[22], and prevents cell apoptosis when overexpressed[23], investigating its role in regulating cortical F-actin in chondrocytes may provide insight into the cytoskeletal mechanisms regulating chondrocyte homeostasis.

Furthermore, elucidating how F-actin regulates gene expression is crucial to understanding the catabolic phenotypic shift of osteoarthritic chondrocytes. Yes-associated protein (YAP) and Myocardin-related transcription factor A (MRTF) are actin-associated transcriptional coactivators expressed in primary chondrocytes[24]. YAP activity is indirectly modulated by F-actin polymerization. Both F-actin and YAP bind to Angiomotin. When F-actin is abundant, Angiomotin binds F-actin. When F-actin depolymerizes, Angiomotin is liberated from F-actin and binds to YAP, sequestering YAP in the cytoplasm[25, 26]. Cytoplasmic YAP no longer binds to TEAD transcription factors, regulating target gene expression. Conversely, G-actin has a high affinity for MRTF. When F-actin depolymerization increases G-actin, G-actin binds to MRTF and sequesters MRTF in the cytoplasm[27, 28]. Cytoplasmic MRTF no longer binds to SRF transcription factors, which regulate target gene expression. Thus, both YAP and MRTF are regulated by F-actin polymerization. In primary chondrocytes, YAP has been shown to regulate *Acan*, *Col2*, and *Sox9* expression[29]. Both YAP and MRTF regulate the expression of *PRG4* expression in primary superficial zone chondrocytes[30, 31]. Additionally, MRTF has been shown to regulate fibroblast and contractile gene expression in passaged chondrocytes[16]. Thus, F-actin loss may regulate the acquisition of catabolic gene programs through YAP and/or MRTF signaling, though this remains to be tested.

In this study, we test the hypotheses that TPM3.1 inhibition will destabilize cortical F-actin, causing dysregulation of chondrocyte homeostasis through YAP and MRTF. These findings will provide new insights into the regulation of osteoarthritis degeneration by actin-mediated signaling.

## Materials and Methods

### Mouse Femoral Cartilage Dissection

Eight-to ten-week-old wild-type (WT), Tpm3.1 knockout (KO, Tpm3/Δexon9d−/−)[32], and knock-in Tpm3.1-neon green (Tpm3.1-NG) mice in the C57BL/6J background (Jackson Laboratories) were gifted from Dr. Edna C. Hardeman and Dr. Peter Gunning (University of New South Wales). Animal procedures were performed in compliance with protocols approved by the University of Delaware Institutional Animal Care and Use Committee.

Femoral heads were dissected from hip joints as described previously[33]. Briefly, following euthanasia, mice were sanitized with 70% (vol/vol) ethanol. An abdominal skin incision was made and retracted to expose the lower torso musculature. The hip joint was dislocated, hind limbs were removed, and surrounding muscle was excised to fully expose the femoral head. Isolated femoral heads were transferred to microcentrifuge tubes containing 1x phosphate-buffered saline (PBS; GenClone; Genesee Scientific, San Diego, CA).

### Culture of Bovine Chondrocytes and Exposure to Pharmacological Inhibitors

Full-thickness chondrocytes were harvested from the cartilage of mature bovine metacarpophalangeal joints, procured from a local butcher, as described previously[7]. Briefly, cells were extracted from cartilage using sequential digestion of 1 hour in 0.5% protease (Sigma-Aldrich; Burlington, MA) in Ham’s F-12 (Corning; Manassas, VA), followed by an overnight digestion in 0.1% collagenase (Roche; Mannheim, Germany) in Ham’s F-12 at 37°C. Digests were filtered, and chondrocytes were counted. Chondrocytes were placed in Ham’s F-12 consisting of 10% FBS (GenClone) and 1% antimycotic/antibiotic (MilliporeSigma; Burlington, MA). Chondrocytes were seeded at 2.5 x 10^5^ cells per well of a 6-well dish or on a glass dish (FluoroDish, FD35–100; World Precision Instruments Inc.; Sarasota, FL) for gene expression and imaging studies, respectively. For protein analysis, 1.875 x 10^6^ cells were seeded within T-75 flasks.

After two days, media was replaced with Ham’s F-12 consisting of 0.5% FBS, 1% antimycotic/antibiotic (serum-reduced media), and pharmacological inhibitors. TPM3.1, YAP, and MRTF were inhibited using TR100 (SML1065, Sigma-Aldrich), verteporfin (HY-B0146, MedChem Express; Monmouth Junction, NJ), and CCG1423 (10010350, Cayman; Ann Arbor, Michigan), respectively, at the concentrations specified in figures. Control groups consisted of chondrocytes treated with Dimethyl Sulfoxide (DMSO; ThermoFisher Scientific; Waltham, MA) vehicle at equal volumes to treatment amount.

### RNA extraction and Relative Real-Time Polymerase Chain Reaction (RT-PCR)

For RNA extraction, cartilage from two femoral heads (one mouse) was scraped off using a scalpel under a brightfield dissection microscope, washed with 1x PBS, placed in 1mL TRIreagent (R2050-1-200, Zymo Research; Irvine, CA), and crushed using a Pellet Pestle. RNA was extracted from cells via chloroform phase separation and purified using the RNA Clean and Concentrator-5 kit (Zymo).

RNA was reverse transcribed into cDNA using UltraScript 2.0 cDNA Synthesis Kit (PCR Biosystems; Wayne, PA). Real-time RT-PCR was performed on equal cDNA concentrations using qPCRBio Sygreen Blue Mix (PCR Biosystems; London, UK) and primers listed in Table S1A. Amplification was performed on a Cielo 3 PCR machine (Azure; Houston, TX). Relative mRNA expression levels were calculated using the ΔΔCT method[34] with 18S as the housekeeping gene. Expression values were normalized to the average of controls of each experiment.

### Sanger Sequencing

Semi-quantitative RT-PCR was performed on equal cDNA concentrations using 2x PCRBIO HS Taq Mix Red (PCR Biosystems; London, UK) and validated Tpm primers[35]. Amplification was performed on a T100 thermocycler (Bio-Rad; Hercules, CA) at 60°C for 40 cycles. PCR products were separated on a 2% agarose gel, and gel bands were cut on a UVP UV Transilluminator (Upland, CA). DNA was purified using the GenCatch Advance Gel Extraction Kit (Epoch Life Sciences; Missouri 167 City, TX) and Sanger sequenced to confirm isoform identity (Genewiz, South Plainfield, NJ).

### WES Capillary Electrophoresis (WES)

G-and F-actin were separated using Triton-solubility[36]. Equal volumes of Triton-soluble (predominantly G-actin) and Triton-insoluble (predominantly F-actin) were used for WES Capillary electrophoresis (Protein Simple; San Jose, CA). Actin was probed using anti-rabbit pan-actin antibody (Table S1B) and secondary-labeled using anti-rabbit HRP-conjugate (Protein Simple). Percent F-actin was calculated by dividing F-actin amount by total actin (sum of G-and F-actin fractions).

To determine specific protein levels, chondrocyte protein was extracted as previously described[8]. Chondrocytes were scraped in radioimmunoprecipitation lysis buffer (Millipore Sigma) with protease inhibitor (Genesee). Cell lysates were sonicated (Model Q55, Qsonica; Newton, CT) and quantified using a bicinchoninic acid assay (Prometheus Protein Biology Products; Genesee). Approximately 0.5-1.0 µg/ml of lysate was separated on 12–230 kDa or 66-440 kDa Separation module kits. Samples were probed using respective primary antibodies (Table S1B), then detected using anti-rabbit HRP-conjugate secondary label (Protein Simple). Specific protein levels were normalized to total protein. Total protein was determined using a Chemiluminescent WES Simple Western Size-Based Total Protein Assay (Protein Simple).

### Immunostaining and Confocal Microscopy

Chondrocytes on glass dishes were fixed with 4% paraformaldehyde at room temperature for 15 minutes, washed in PBS, and placed in blocking/permeabilization buffer (3% Goat serum, 3% BSA, and 0.3% Triton in 1xPBS) for 15 min. Chondrocytes were incubated with primary antibodies against YAP or MRTF (Table S1B), then stained in a permeabilization/blocking buffer solution containing Alexa 488 Rabbit (1:500, Biotium; Fremont, CA), Rhodamine Phalloidin (1:50, Biotium), and Hoechst (1:500, Biotium). Chondrocytes were washed, coverslip-mounted with Drop-n-Stain EverBrite™ Mounting Medium (Biotium), and imaged on a Zeiss LSM880 confocal microscope using 0.3µm z-stack steps. Unless otherwise specified, a 63X objective was used.

For whole-mount imaging, mouse femoral heads were fixed in 4% paraformaldehyde at 4°C overnight, placed in permeabilization-blocking buffer for 30 minutes, then stained in permeabilization/blocking buffer containing Rhodamine Phalloidin (1:20) and Hoechst (1:500). Femoral heads were mounted *en face* in a glass well dish within an agarose divot using an adapted whole-mount imaging protocol[37]. Zeiss LSM880 confocal microscope equipped with a 40x objective, 0.3µm z-stack step, at 1X zoom.

Images were processed using ZEN (Zeiss, Black Edition) and analyzed with FIJI ImageJ. Single Z-stack slices are shown from chondrocyte lateral regions displaying cortical F-actin. For image quantification, Rhodamine Phalloidin was used as a cell border indicator, and cells were traced manually as previously described[8]. Mean fluorescence for F-actin, YAP, or MRTF was then quantified.

### Statistics

All experiments were conducted on separate occasions with samples from ≥3 separate animals. Statistical analysis was performed using GraphPad Prism 11 (San Diego, CA). Experiments were normalized to controls and pooled together for statistics. Unpaired t-tests were used to detect differences between two data groups. An analysis of variance (ANOVA) followed by a Dunnett post hoc test was used to determine the differences between three or more data groups. Outliers were identified using the ROUT method with a maximum desired false discovery rate (Q) set at 1%[38]. All graphs are represented as averages ± standard error of the mean.

## Results

### Identification of Tpm isoform expression

To identify the Tpm isoforms expressed in chondrocytes, we isolated mouse femoral head cartilage and extracted chondrocyte RNA. We performed semi-quantitative RT-PCR using Tpm primers[35] followed by gel electrophoresis of cDNA. Chondrocytes express seven Tpm isoforms (Figure 1A). Since closely related Tpm isoforms can share terminal exon 1 and exon 9, we performed Sanger sequencing to identify internal exons (Figure S1). Sanger sequencing revealed the expression of *Tpm1.6*, *1.8*, *1.13*, *2.1*, *2.2*, *3.1*, and *4.2* in native mouse chondrocytes. Consequently, *Tpm1.2*, *1.7*, *1.9*, *2.3*, and *3.2* are likely not expressed by chondrocytes due to poor sequence alignments with these isoforms. Additionally, chondrocytes did not have visible bands for *Tpm1.1*, *1.5*, *1.10*, and *1.12,* suggesting unlikely expression for these Tpm isoforms.

**Figure 1.**
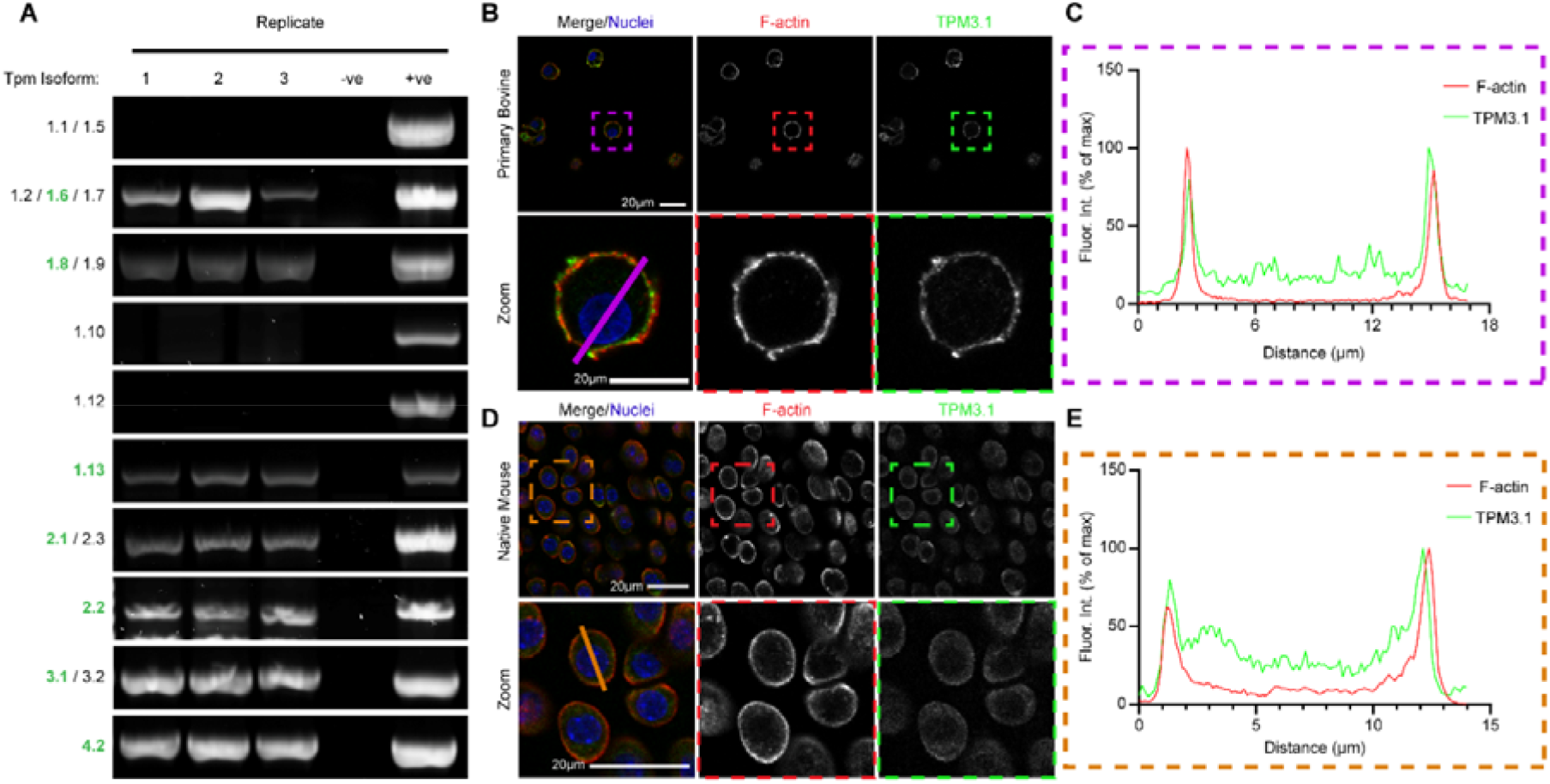
Tropomyosin 3.1 Expression and Localization in Chondrocytes. (A) DNA gel electrophoresis followed by Sanger sequencing demonstrates the tropomyosin isoforms expressed in mouse chondrocytes. (B) Isolated primary bovine chondrocytes showing colocalization of F-actin and Tropomyosin 3.1 (Red = phalloidin for F-actin; Blue = Hoechst for nuclei; Green = Alexa 488 for Tropomyosin 3.1). (C) Fluorescence intensity and localization of F-actin and Tropomyosin 3.1 along the purple line in B. (D) Native mouse femoral head chondrocytes showing colocalization of F-actin and Tropomyosin 3.1 (Red = phalloidin for F-actin; Blue = Hoechst for nuclei; Green = Alexa 488 for Tropomyosin 3.1). (E) Fluorescence intensity and localization of F-actin and Tropomyosin 3.1 along the orange line in D (n = 3 mice, 1 femoral head from each).

Previously, we and others have shown that TPM3.1 regulates F-actin stress fiber stability in passaged chondrocytes[8]. However, the role TPM3.1 plays in regulating cortical F-actin in primary chondrocytes remains elusive. We investigated TPM3.1 localization and observed that TPM3.1 colocalizes with cortical F-actin in primary bovine chondrocytes (Figure 1B). A line scan through the chondrocyte revealed overlapping peaks in intensity for TPM3.1 and F-actin, demonstrating association along the cortex (Figure 1C), consistent with previous work[8]. We also determined F-actin and TPM3.1 colocalization *ex vivo* in mouse femoral head cartilage. Using TPM3.1-NG mice that express an mNeonGreen (NG) tag in the C-terminus of Tpm3.1[39], we found that TPM3.1 associates with cortical F-actin (Figure 1D). Consistent with primary bovine chondrocytes, a line scan through the cortex of native mouse chondrocytes revealed overlapping peaks in intensity for TPM3.1 and F-actin on the cortex (Figure 1E).

### TPM3.1 regulates chondrocyte cortical F-actin

To determine cortical F-actin regulation by TPM3.1, we examined cortical F-actin organization in native femoral head chondrocytes from Tpm3.1-/-mice[32]. As compared to Tpm3.1+/+ chondrocytes, Tpm3.1-/- chondrocytes had reduced F-actin staining. On average, F-actin fluorescence intensity in Tpm3.1-/- chondrocytes was 60.2% ± 7.3% (average ± SEM) of Tpm3.1+/+ chondrocytes (Figure S2).

Next, we examined the effect of TPM3.1 inhibitor TR100 on native chondrocyte F-actin. TR100 inhibits TPM3.1 by binding to the C-terminus (encoded by the 9d exon) of free TPM3.1. This compromises TPM3.1 filament integrity along F-actin[40], contributing to F-actin instability. We found that chondrocytes exposed to ≤4µM TR100 are viable (Figure S3). Native cartilage exposed to TR100 alters F-actin and TPM3.1 localization along the cortex as seen via line scan analysis (Figure 2B).

**Figure 2.**
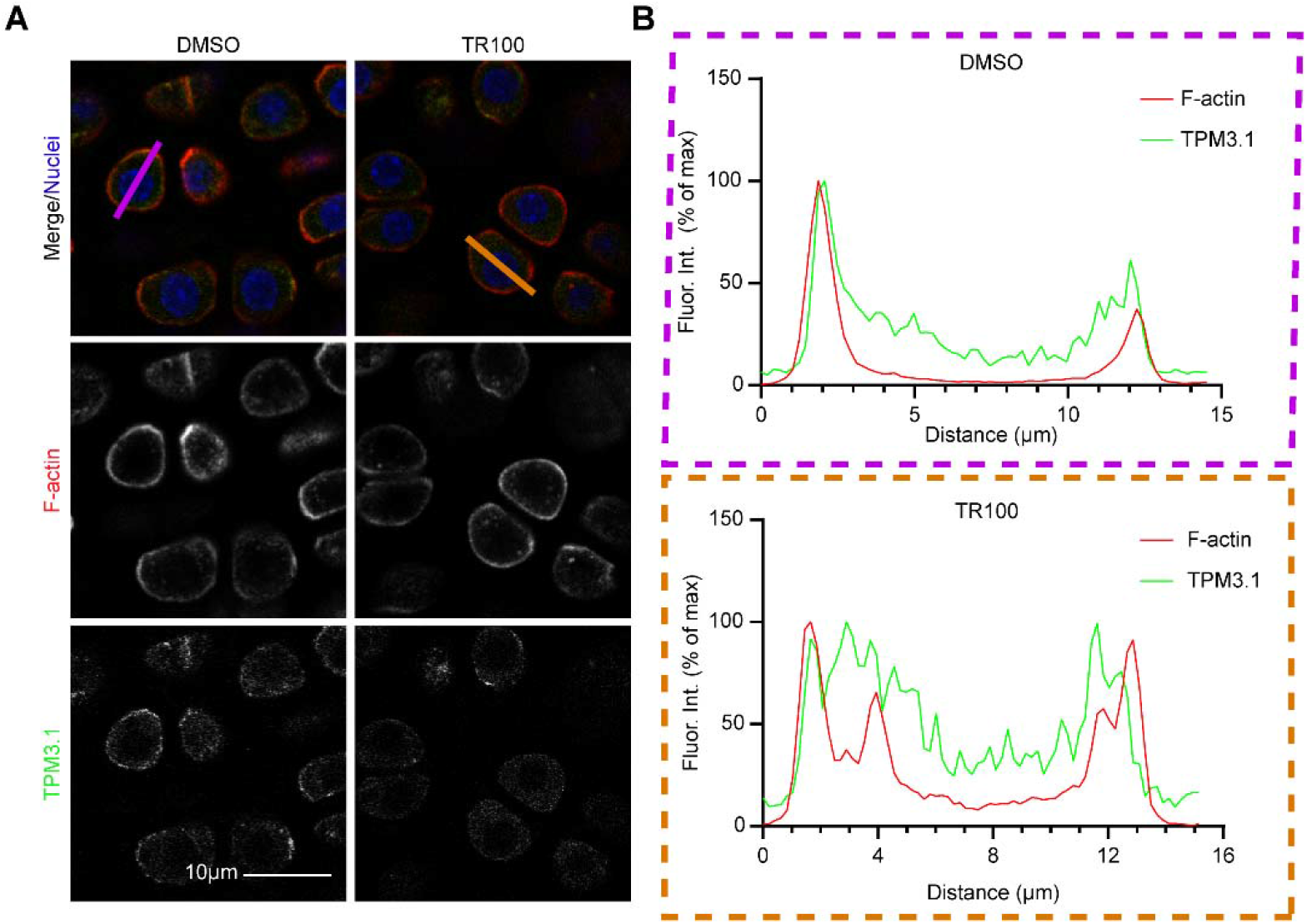
TR100 Treatment Dysregulates Colocalization of Tropomyosin 3.1 and F-actin. (A) Organization of F-actin and Tropomyosin 3.1 in mouse femoral head chondrocytes treated with DMSO control, left, and TR100, right for 24 hours (Red = phalloidin for F-actin; Blue = Hoechst for nuclei; Green = Alexa 488 for Tropomyosin 3.1). (B) Fluorescence intensity of F-actin and Tropomyosin 3.1, showing colocalization in DMSO-treated chondrocytes (purple line/box) and dysregulated organization in TR100-treated chondrocytes (orange line/box) (n = 3 mice, left hip DMSO, right hip TR100 treated).

Furthermore, TR100 decreased F-actin fluorescent intensity in bovine chondrocytes to 60.2% ± 6.6% and 41.4% ± 4.0% of controls for 2 and 4μM TR100, respectively (Figure 3A, B). Additionally, using WES capillary electrophoresis (WES), we found that TR100 reduced the total actin in chondrocytes to 32.7% ± 9.2% of controls (Figure 3C, D). Additionally, WES analysis confirmed that TR100 reduced the proportion of F-actin (Figure 3E, F) in chondrocytes.

**Figure 3.**
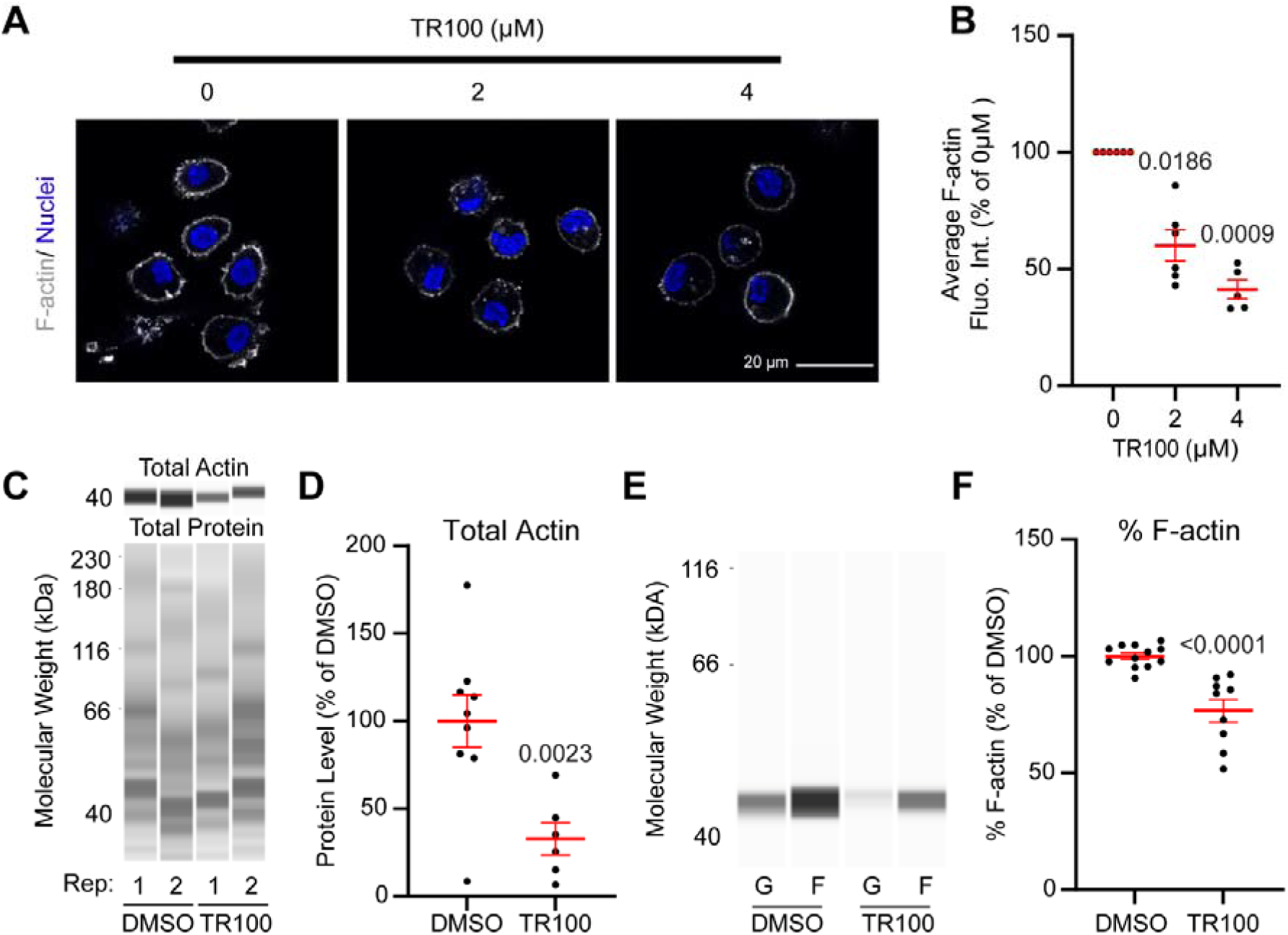
Tropomyosin 3.1 Inhibition via TR100 Treatment Reduces Cortical F-actin. (A) Confocal microscopy of TR100-treated chondrocytes after 24 hours (Grey = phalloidin for F-actin; Blue = Hoechst for nuclei). (B) Dot plot of TR100 treatment measuring F-actin fluorescence intensity as a percent of the control; each point represents the average of 10-15 chondrocytes per condition (n=6 trials). (C) Wes Capillary Electrophoresis (WES) pseudo-blot and (D) quantification of changes in total actin normalized to total protein of DMSO-and TR100-treated cells after 48 hours of treatment; each trial was made up of three replicates and normalized to controls (n= 3 trials). (E) WES pseudo-blot and (F) quantification of changes in percent soluble G-actin (G) and insoluble F-actin (F) in TR100-treated cells treated for 24 hours normalized to DMSO control; each trial was made up of three replicates and normalized to controls (n= 3 trials).

### TPM3.1 inhibition using pharmacological agent, TR100, dysregulates chondrocyte homeostasis

While stress fiber F-actin loss through TPM3.1 inhibition is known to regulate the phenotype of dedifferentiated, passaged chondrocytes[8], we aimed to determine whether TPM3.1 inhibition alters homeostasis in primary chondrocytes. We found that TR100 treatment decreased cartilage mRNA levels of *ACAN* to 56.7± 7.9%*, COL2* to 11.1% ± 2.6%, and *PRG4* to 26.6% ± 4.8% of untreated controls. In addition, TR100 treatment increased degradative enzyme *MMP3* mRNA levels to 972.2% ± 203.8% of control levels (Figure 4). TR100 treatment did not affect *MMP13* mRNA levels.

**Figure 4.**
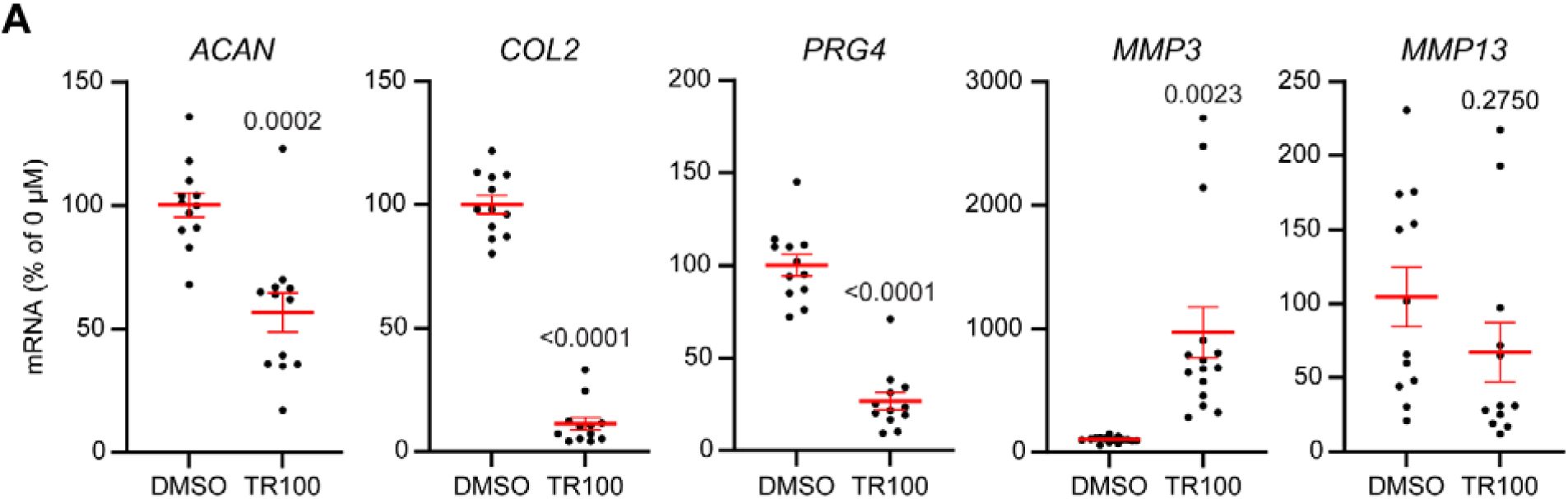
Effects of TPM3.1 Inhibition on Chondrocyte Homeostasis. RT-PCR analysis of primary bovine chondrocytes treated with DMSO or 4μM of TR100 for 24 hours. Chondrogenic genes Aggrecan (*ACAN*), Collagen II (*COL2*), lubricative molecule Proteoglycan 4 (*PRG4*), and degenerative enzymes Matrix Metalloproteinases (*MMP*) -3 and -13 were used as markers of change in chondrocyte homeostasis; each trial was made up of three replicates and normalized to controls (n= 4-5 trials).

### F-actin destabilization regulates chondrocyte phenotype by regulating YAP signaling

Since F-actin loss by TPM3.1 inhibition shifted primary chondrocyte mRNA levels towards catabolism, we next investigated the downstream regulation of gene expression by cortical F-actin destabilization. Alterations in F-actin organization can mediate gene expression directly through actin-binding transcription factors, such as YAP and MRTF[29–31]. YAP is a transcriptional co-activator whose nuclear localization is promoted by a high proportion of F-actin[26]; thus, F-actin depolymerization is expected to reduce nuclear YAP, altering expression of YAP-mediated genes. We found that TR100 reduces the nuclear-to-cytoplasmic YAP proportion to 72.6% ± 6.2% and 49.9% ± 3.0% of controls for 2mM and 4µM TR100 (Figure 5A, B). To further elucidate the contribution of YAP-mediated changes in gene expression, we treated chondrocytes with YAP inhibitor, verteporfin. Chondrocytes exposed to ≤5µM verteporfin are viable (Figure S4). We determined that verteporfin treatment of primary chondrocytes does not alter the nuclear-to-cytoplasmic YAP ratio as compared to controls (Figure 5C, D). This may be expected, as verteporfin disrupts the binding of YAP to TEAD[41] to prevent associated gene transcription. YAP can still enter the nucleus following verteporfin treatment. Nevertheless, we found verteporfin treatment decreases *COL2* and *PRG4* mRNA levels to 77.2% ± 7.6% and 57.3% ± 10.0% of controls, respectively. *ACAN* or *MMP3* mRNA levels were unchanged following verteporfin treatment (Figure 5E).

**Figure 5.**
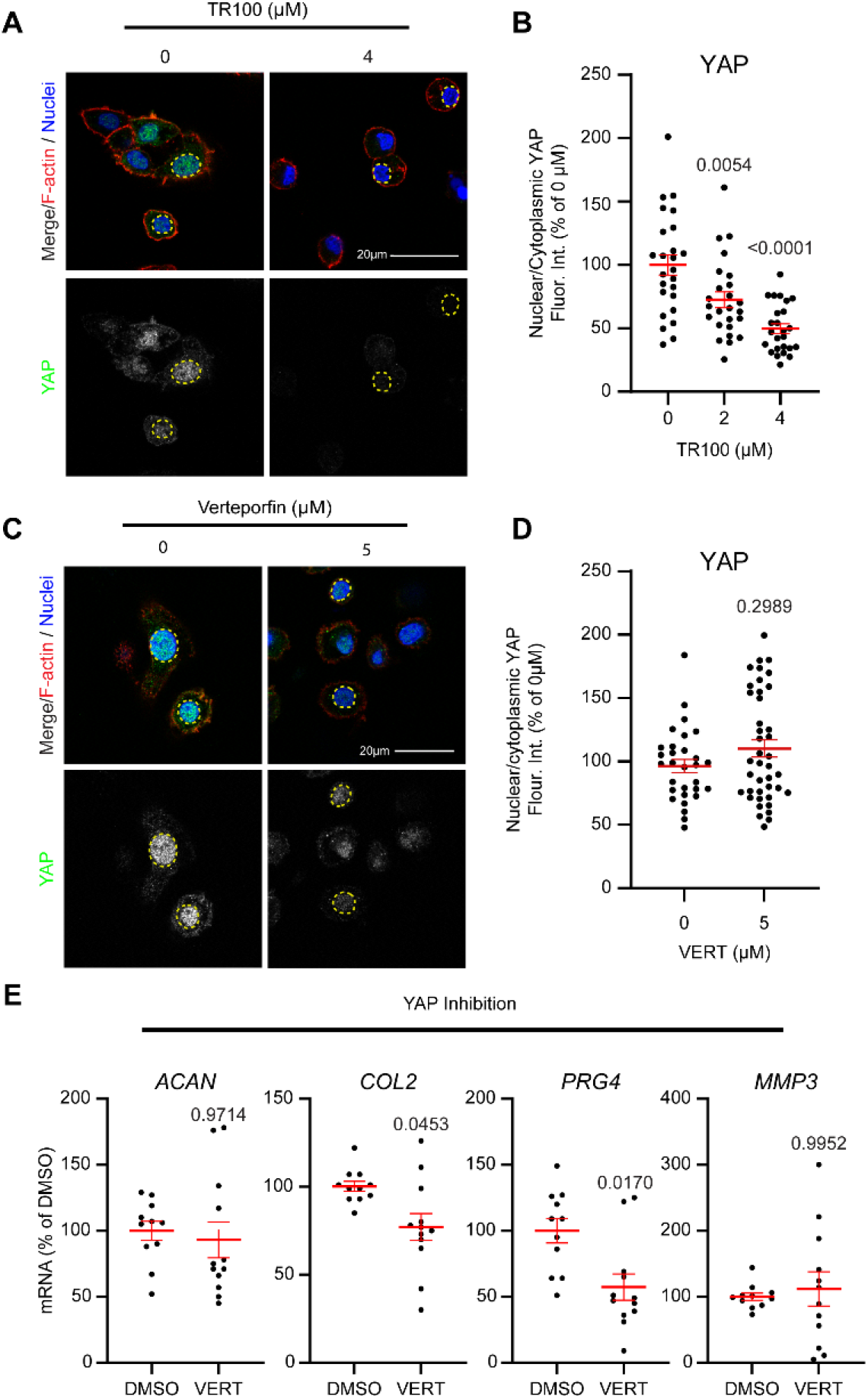
mRNA Regulation by YAP/TAZ Signaling. (A) Chondrocytes treated with 4µM TR100 for 24 hours and stained for YAP to examine alterations in nuclear localization (Red = phalloidin for F-actin; Blue = Hoechst for nuclei; Green = Alexa 488 for YAP). (B) Quantification of nuclear-to-cytoplasmic ratio of YAP fluorescence intensity in TR100-treated chondrocytes; 10-15 chondrocytes used per trial, normalized to average DMSO control (n=3 trials). (C) Confocal microscopy of chondrocytes treated with 5µM of YAP inhibitor Verteporfin (Vert) for 24 hours, visualizing nuclear vs cytoplasmic localization of YAP (Red = phalloidin for F-actin; Blue = Hoechst for nuclei; Green = Alexa 488 for YAP). (D) Quantification of nuclear-to-cytoplasmic ratio of YAP fluorescence intensity in Vert-treated chondrocytes; 10-15 chondrocytes used per trial, normalized to average DMSO control (n=3 trials). (E) RT-PCR of chondrocytes treated with 5µM Vert for 24 hours. Chondrogenic genes *ACAN*, *COL2*, and *PRG4*, and degradative enzyme *MMP3* were used as markers for alterations in chondrocyte homeostasis; each trial was made up of three replicates and normalized to controls (n= 3-4 trials).

### F-actin destabilization regulates chondrocyte phenotype by regulating MRTF signaling

Similar to YAP, MRTF is a downstream actin-mediated transcriptional co-activator found to be expressed in primary chondrocytes[24], whose nuclear localization is regulated by actin polymerization[27, 28]. MRTF has a high affinity for G-actin; therefore, F-actin depolymerization reduces nuclear MRTF[36, 42]. In primary chondrocytes, exposure to TR100 decreases the MRTF nuclear-to-cytoplasmic ratio to 88.6% ± 3.8% and 82.6% ± 2.7% for 2mM and 4µM TR100 relative to controls (Figure 6A, B). Next, we treated chondrocytes with the MRTF inhibitor, CCG1423, which inhibits the MRTF nuclear import[43]. Chondrocytes treated with ≤10µM of CCG1423 are viable (Figure S4). CCG1423 reduces the MRTF nuclear-to-cytoplasmic ratio to 69.9% ± 1.6% of controls (Figure 6 C, D). Furthermore, CCG1423 treatment decreases *ACAN*, *COL2*, *PRG4* and *MMP3* mRNA levels to 73.7% ± 8.1%, 35.1% ± 9.2%, 66.0% ± 10.7% and 40.1% ± 12.6% of control mRNA levels, respectively (Figure 6E).

**Figure 6.**
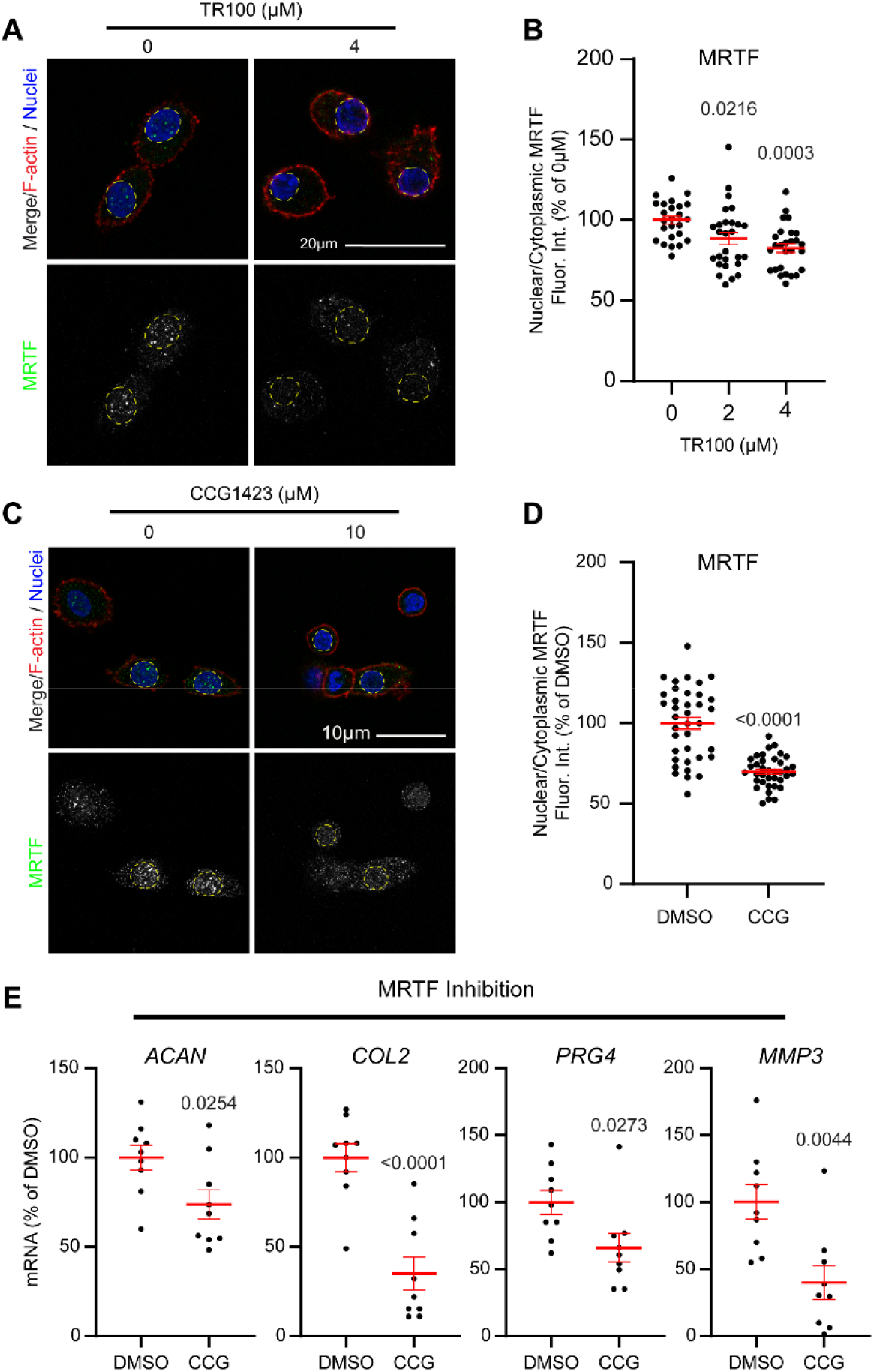
mRNA Regulation by MRTF Signaling. (A) Chondrocytes treated with 4µM TR100 for 24 hours and stained for MRTF to examine alterations in nuclear localization (Red = phalloidin for F-actin; Blue = Hoechst for nuclei; Green = Alexa 488 for MRTF). (B) Quantification of nuclear-to-cytoplasmic ratio of MRTF fluorescence intensity in TR100-treated chondrocytes; 10-15 chondrocytes used per trial, normalized to average DMSO control (n=3 trials). (C) Confocal microscopy of chondrocytes treated with 10µM of MRTF inhibitor CCG1423 (CCG) for 24 hours, visualizing nuclear vs cytoplasmic localization of MRTF (Red = phalloidin for F-actin; Blue = Hoechst for nuclei; Green = Alexa 488 for MRTF). (D) Quantification of nuclear-to-cytoplasmic ratio of MRTF fluorescence intensity in CCG-treated chondrocytes; 10-15 chondrocytes used per trial, normalized to average DMSO control (n=3 trials). (E) RT-PCR of chondrocytes treated with CCG for 24 hours. Chondrogenic genes *ACAN*, *COL2*, and PRG*4*, and degradative enzyme *MMP3* were used as markers for alterations in chondrocyte homeostasis; each trial was made up of three replicates and normalized to controls (n=3 trials).

### TPM3.1 inhibition dysregulates COL2 and MMP3 protein levels

Our findings above suggest cortical F-actin loss regulates gene expression at least partially through YAP and MRTF. To determine if cortical F-actin loss also altered protein levels, we examined the effects of TR100 on cartilage matrix molecule, COL2, and degradative enzyme, MMP3, protein levels. We found that, on average, TR100 reduces COL2 protein levels to 11.4% ± 7.4% of DMSO controls (Figure 7A, B), whereas TR100 increases MMP3 protein levels to 573.3% ± 52.4% of DMSO controls (Figure 7C, D).

**Figure 7.**
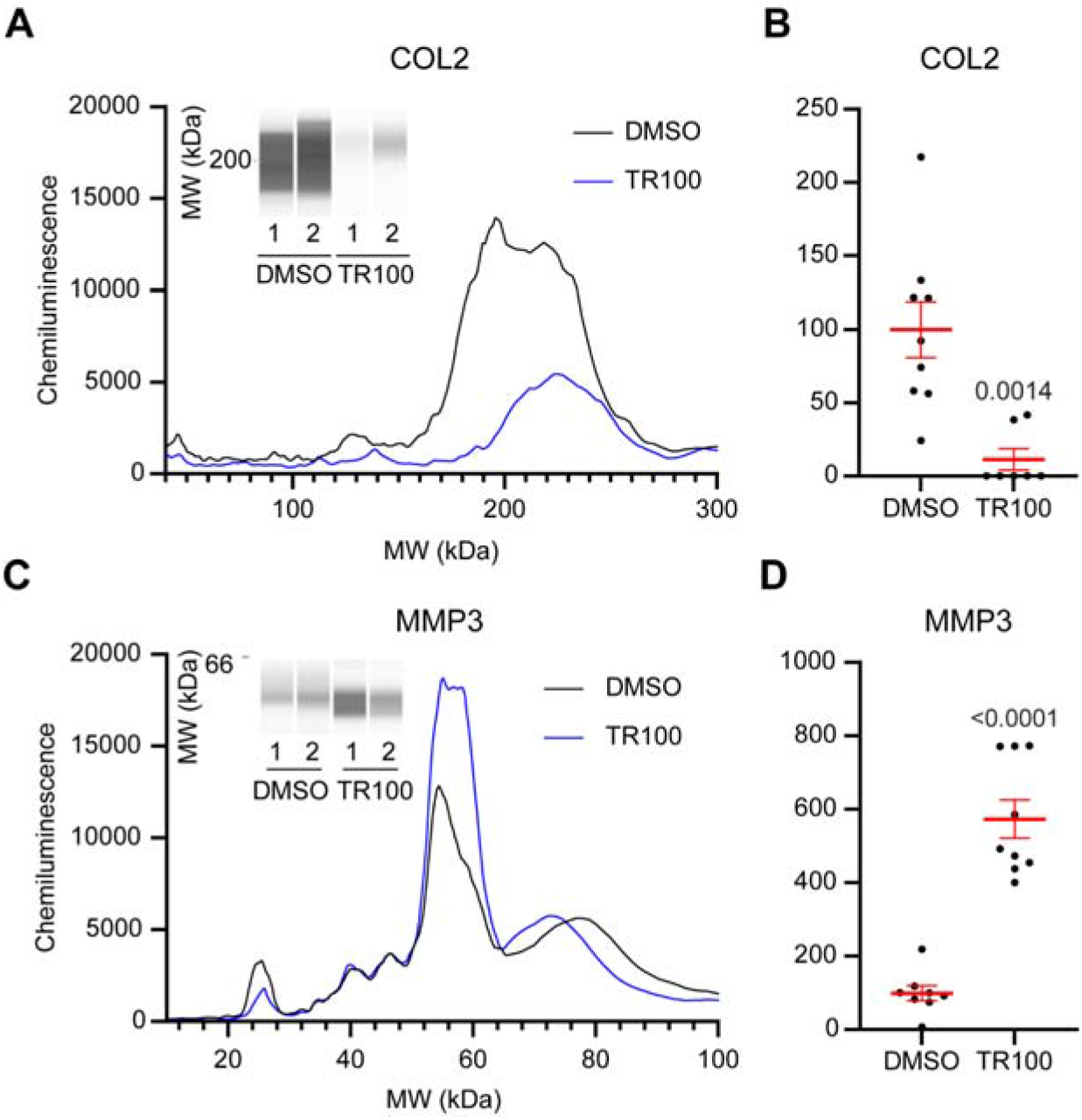
TPM3.1 Inhibition Causes a Catabolic Shift in Chondrocyte Homeostasis. WES analysis of primary bovine chondrocytes treated with TR100 for 48 hours. (A) COL2 WES pseudo-blot and electropherogram demonstrating chemiluminescence for DMSO-and TR100-treated samples. (B) Dot plot of COL2; each trial was made up of three replicates and normalized to controls (n=3 trials). (C) MMP3 WES pseudo-blot and electropherogram demonstrating chemiluminescence for DMSO-and TR100-treated samples. (D) Dot plot for MMP3; each trial was made up of three replicates and normalized to controls (n=3 trials).

## Discussion

These findings provide a greater understanding of the role cortical F-actin stability plays in regulating chondrocyte homeostasis. We hypothesized that cortical F-actin destabilization by TPM3.1 perturbation would dysregulate chondrocyte homeostasis by a reduction in cortical F-actin and alteration of downstream actin-mediated signaling pathways. Our data lend support for our hypothesis. TPM3.1 perturbation, either through *Tpm3.1* knockout or TPM3.1 inhibition, reduces cortical F-actin in both native and primary chondrocytes. Furthermore, TPM3.1 inhibition in primary chondrocytes results in a catabolic shift. The reduction in cortical F-actin caused by TPM3.1 inhibition caused nuclear export of actin-mediated transcription factors, YAP and MRTF, which partially regulate the catabolic shift. Collectively, these findings identify TPM3.1 as a regulator of cortical F-actin stability and establish downstream mechanisms by which cortical F-actin levels regulate chondrocyte homeostasis.

The TPM isoform, TPM3.1, regulates cortical F-actin stability. The expression of Tpm3.1 in mouse and bovine primary chondrocytes agrees with our previous work in human chondrocytes[8], demonstrating cross-species expression of Tpm3.1 in chondrocytes. TPM3.1 has been shown previously to associate with F-actin and regulate stability in other cell types[20, 44]. Our previous study focused on the role that TPM3.1 plays in regulating the stability of F-actin stress fibers in passaged chondrocytes[8]; however, our new findings demonstrate that TPM3.1 also regulates cortical F-actin in native and primary chondrocytes. Cortical F-actin regulation by TPM3.1 is consistent with findings in B-cells and white adipose tissue, which showed that treatment with anti-TPM drugs or Tpm3.1 knockout reduces cortical F-actin[45, 46]. While we found that Tpm3.1 loss reduces cortical F-actin, cortical F-actin remains present in Tpm3.1-deficient chondrocytes. In addition to TPM3.1, we found that chondrocytes express at least six other Tpm isoforms: 1.6, 1.8, 1.13, 2.1, 2.2, and 4.2. While it is unclear whether these other isoforms regulate chondrocyte cortical actin, studies in breast epithelial cells have demonstrated that TPM2.1 knockdown may disrupt cortical F-actin[47]. Therefore, Tpm3.1 knockout may result in functional compensation from other tropomyosin isoforms, such as Tpm2.1. This redundancy may be particularly important in chondrocytes, where cortical F-actin preservation appears critical for maintaining chondrocyte phenotype. Nevertheless, Tpm3.1 may not be required for cortical actin formation. However, since its loss reduces cortical F-actin, TPM3.1 appears important for maintaining high cortical F-actin levels. In support of this, in primary chondrocytes that express Tpm3.1, we found that TPM3.1 inhibition via TR100 mediated cortical F-actin loss. This resulted in a catabolic shift in chondrocyte gene expression.

Our study demonstrates cortical F-actin as a critical regulator of chondrocyte homeostasis. This corroborates findings from other studies where cortical F-actin loss correlates with cartilage degeneration[12, 13]. Previously, we found decreased cortical F-actin following destabilization of the medial meniscus, coinciding with cartilage degeneration[12]. Further support from human cartilage shows reduced levels of staining for cortical F-actin in chondrocytes from osteoarthritic cartilage as compared to healthy cartilage[13]. Our current study extends these findings and provides new insights into the causal relationship between F-actin loss and degeneration. Cortical F-actin reduction, through inhibiting TPM3.1, causes a catabolic shift in the regulation of critical chondrogenic and degradative genes. Intriguingly, the perturbation of TPM3.1 was unfavorable for the primary chondrocyte phenotype, which is in contrast to the favorable effect of TPM3.1 inhibition in passaged chondrocytes[8]. We previously found TR100 treatment to be beneficial in passaged chondrocytes by repressing the dedifferentiated phenotype. These contrasting responses highlight that chondrocyte homeostasis depends not solely on the F-actin amount, but on its appropriate organization. Maintenance of F-actin as a cortical network appears critical for the differentiated chondrocyte phenotype, as cortical F-actin reduction promotes a catabolic phenotype, whereas stress fiber formation is associated with chondrocyte dedifferentiation. While the findings of the present study could suggest that enhancing cortical F-actin stability via TPM3.1 may provide an attractive route to prevent osteoarthritis progression, it is unclear if TPM3.1 enhancement would instead promote stress fiber formation and subsequent dedifferentiation. Therefore, an understanding of the mechanisms that determine whether chondrocytes present cortical versus stress fiber F-actin organization is needed.

To define a mechanism by which cortical F-actin stability regulates chondrocyte homeostasis, we examined the actin-responsive transcriptional coactivators YAP and MRTF. In chondrocytes, these pathways have largely been studied in the context of superficial zone phenotype and chondrocyte dedifferentiation[16, 24, 48]. Here, we found that TPM3.1 inhibition reduced YAP and MRTF nuclear localization. Furthermore, direct pharmacological inhibition of either YAP or MRTF altered chondrocyte gene expression. In keeping with previous studies in superficial zone chondrocytes, inhibition of either YAP or MRTF reduced *PRG4* expression, supporting roles in regulating the superficial zone phenotype[30, 48]. Importantly, MRTF and YAP contribute more broadly to chondrocyte homeostasis. We found that inhibition of these pathways altered additional cartilage matrix genes, including *ACAN* and *COL2*, extending their potential roles beyond *PRG4* regulation. Our findings are also in agreement with previous YAP inhibition studies, which found that Verteporfin treatment led to *ACAN*, *COL2*, and *SOX9* loss in primary chondrocytes[29]. Together, these findings support YAP and MRTF as downstream mechanisms through which cortical F-actin stability is translated into changes in specific chondrocyte gene expression. Interestingly, *MMP3* regulation differed from cartilage matrix genes. Although TPM3.1 inhibition increased *MMP3* expression, YAP inhibition did not affect *MMP3*, whereas MRTF inhibition downregulated *MMP3*. This dichotomy suggests divergent pathways regulate *MMP3*. The increase in *MMP3* expression following cortical F-actin destabilization cannot be readily explained by reduced YAP or MRTF activity and may instead involve additional actin-sensitive transcriptional mechanisms. One potential mechanism involves cysteine-rich protein 2 (CRP2), an F-actin-binding protein identified as a link between actin depolymerization and MMP expression. In breast cancer cells, F-actin depolymerization promotes CRP2 translocation to the nucleus, where CRP2 regulates expression of several MMPs (i.e., *MMP9* and *MMP13*)[17]. Whether a similar actin-CRP2 signaling mechanism regulates *MMP3* in chondrocytes remains unknown. Nevertheless, these findings demonstrate cortical F-actin loss regulates chondrocyte homeostasis through several distinct actin-sensitive transcriptional pathways.

While our findings provide insight into the downstream mechanisms by which actin regulates chondrocyte homeostasis, one unanswered question is how TPM3.1-mediated cortical F-actin stability is regulated in osteoarthritis. In post-traumatic osteoarthritis, cortical F-actin loss may be due to mechanical overloading. Previous studies demonstrated that cyclic compression reduces cortical F-actin in chondrocytes and promotes actin remodeling, which was suggested to be due to actin-severing (depolymerization) protein activity[49]. Intriguingly, TPM3.1 has been shown to prevent the interaction between actin-severing protein, cofilin, and F-actin[50]. We postulate that mechanical overload may reduce the interaction between TPM3.1 and F-actin, resulting in F-actin exposure to severing proteins. Upstream mechanosensitive pathways, including Rho-family GTPases, represent potential regulators of this process. Previously, we found CDC42 inhibition reduced the association of TPM3.1 with F-actin in passaged chondrocytes[8]. Therefore, it may be plausible that load reduces CDC42 activity and thereby reduces TPM3.1-F-actin association. Cortical F-actin loss during joint injury may represent an actively regulated response to altered mechanical loading rather than mechanical breakage of F-actin. Determining whether mechanical overload alters TPM3.1 localization or association with cortical F-actin will therefore be important for understanding how abnormal loading contributes to cytoskeletal disruption during cartilage degeneration.

In summary, our findings establish cortical F-actin stability as a chondrocyte homeostasis regulator and TPM3.1 as a cortical F-actin stabilizer. These findings support a model in which cortical actin cytoskeleton disruption is not simply a consequence of cartilage degeneration but instead contributes to the phenotypic changes driving osteoarthritis progression. Importantly, our findings also demonstrate that the consequences of altering F-actin are dependent on its organization, emphasizing that cortical F-actin preservation, rather than increasing or decreasing F-actin polymerization, is critical for maintaining the differentiated chondrocyte phenotype. Further defining the mechanisms that regulate TPM3.1 and cortical F-actin organization in response to mechanical loading may therefore reveal new approaches for maintaining chondrocyte homeostasis and limiting degeneration in osteoarthritis.

## Acknowledgements

The authors would like to acknowledge Audrey Houghton for her assistance in quantification.

## Author Contribution

Marin Herrick: investigation, data curation, formal analysis, visualization, writing-original draft. Mark Arranguez: investigation, data curation, formal analysis, visualization, writing-review and editing. Josalyn Stack: formal analysis, writing-review and editing. Justin Parreno: Conceptualization, resources, funding acquisition, supervision, writing-review and editing.

## Use of AI

Artificial intelligence technologies were NOT used in the collection or processing of data, NOR was it used in the writing and editing process of this manuscript.

## References

1. Aigner, T., et al., Roles of chondrocytes in the pathogenesis of osteoarthritis. Curr Opin Rheumatol, 2002. 14(5): p. 578–84.

2. Saarakkala, S., et al., Depth-wise progression of osteoarthritis in human articular cartilage: investigation of composition, structure and biomechanics. Osteoarthritis Cartilage, 2010. 18(1): p. 73–81.

3. Rizkalla, G., et al., Studies of the articular cartilage proteoglycan aggrecan in health and osteoarthritis. Evidence for molecular heterogeneity and extensive molecular changes in disease. J Clin Invest, 1992. 90(6): p. 2268–77.

4. Dodge, G.R. and A.R. Poole, Immunohistochemical detection and immunochemical analysis of type II collagen degradation in human normal, rheumatoid, and osteoarthritic articular cartilages and in explants of bovine articular cartilage cultured with interleukin 1. J Clin Invest, 1989. 83(2): p. 647–61.

5. Mitchell, P.G., et al., Cloning, expression, and type II collagenolytic activity of matrix metalloproteinase-13 from human osteoarthritic cartilage. J Clin Invest, 1996. 97(3): p. 761–8.

6. Chubinskaya, S., K.E. Kuettner, and A.A. Cole, Expression of matrix metalloproteinases in normal and damaged articular cartilage from human knee and ankle joints. Lab Invest, 1999. 79(12): p. 1669–77.

7. Parreno, J., et al., Interplay between cytoskeletal polymerization and the chondrogenic phenotype in chondrocytes passaged in monolayer culture. J Anat, 2017. 230(2): p. 234–248.

8. Schofield, M.M., et al., Targeting F-actin stress fibers to suppress the dedifferentiated phenotype in chondrocytes. Eur J Cell Biol, 2024. 103(2): p. 151424.

9. Lauer, J.C., et al., Articular Chondrocyte Phenotype Regulation through the Cytoskeleton and the Signaling Processes That Originate from or Converge on the Cytoskeleton: Towards a Novel Understanding of the Intersection between Actin Dynamics and Chondrogenic Function. Int J Mol Sci, 2021. 22(6).

10. Mallein-Gerin, F., R. Garrone, and M. van der Rest, Proteoglycan and collagen synthesis are correlated with actin organization in dedifferentiating chondrocytes. Eur J Cell Biol, 1991. 56(2): p. 364–73.

11. Langelier, E., et al., The chondrocyte cytoskeleton in mature articular cartilage: structure and distribution of actin, tubulin, and vimentin filaments. J Histochem Cytochem, 2000. 48(10): p. 1307–20.

12. Chan, B., et al., Adseverin, an actin-binding protein, modulates hypertrophic chondrocyte differentiation and osteoarthritis progression. Sci Adv, 2023. 9(31): p. eadf1130.

13. Lee, W., et al., Inflammatory signaling sensitizes Piezo1 mechanotransduction in articular chondrocytes as a pathogenic feed-forward mechanism in osteoarthritis. Proc Natl Acad Sci U S A, 2021. 118(13).

14. Capin-Gutierrez, N., et al., Cytoskeleton disruption in chondrocytes from a rat osteoarthrosic (OA) -induced model: its potential role in OA pathogenesis. Histol Histopathol, 2004. 19(4): p. 1125–32.

15. Pascarelli, N.A., et al., Changes in Ultrastructure and Cytoskeletal Aspects of Human Normal and Osteoarthritic Chondrocytes Exposed to Interleukin-1beta and Cyclical Hydrostatic Pressure. Int J Mol Sci, 2015. 16(11): p. 26019–34.

16. Parreno, J., et al., Expression of type I collagen and tenascin C is regulated by actin polymerization through MRTF in dedifferentiated chondrocytes. FEBS Lett, 2014. 588(20): p. 3677–84.

17. Mgrditchian, T., et al., Actin cytoskeleton depolymerization increases matrix metalloproteinase gene expression in breast cancer cells by promoting translocation of cysteine-rich protein 2 to the nucleus. Front Cell Dev Biol, 2023. 11: p. 1100938.

18. Sanka, K., et al., Influence of actin cytoskeletal integrity on matrix metalloproteinase-2 activation in cultured human trabecular meshwork cells. Invest Ophthalmol Vis Sci, 2007. 48(5): p. 2105–14.

19. Hardeman, E.C., N.S. Bryce, and P.W. Gunning, Impact of the actin cytoskeleton on cell development and function mediated via tropomyosin isoforms. Semin Cell Dev Biol, 2020. 102: p. 122–131.

20. Bedi, A., et al., The tropomyosin 3.1/3.2 inhibitor ATM-3507 alters B-cell actin dynamics and impairs the growth and motility of diffuse large B-cell lymphoma cell lines. Front Immunol, 2025. 16: p. 1668379.

21. Sui, Z., et al., Stabilization of F-actin by tropomyosin isoforms regulates the morphology and mechanical behavior of red blood cells. Mol Biol Cell, 2017. 28(19): p. 2531–2542.

22. Jalilian, I., et al., Cell elasticity is regulated by the tropomyosin isoform composition of the actin cytoskeleton. PLoS One, 2015. 10(5): p. e0126214.

23. Desouza-Armstrong, M., P.W. Gunning, and J.R. Stehn, Tumor suppressor tropomyosin Tpm2.1 regulates sensitivity to apoptosis beyond anoikis characterized by changes in the levels of intrinsic apoptosis proteins. Cytoskeleton (Hoboken), 2017. 74(6): p. 233–248.

24. Parreno, J., et al., MRTF-A signaling regulates the acquisition of the contractile phenotype in dedifferentiated chondrocytes. Matrix Biol, 2017. 62: p. 3–14.

25. Mana-Capelli, S., et al., Angiomotins link F-actin architecture to Hippo pathway signaling. Mol Biol Cell, 2014. 25(10): p. 1676–85.

26. Das, A., et al., YAP Nuclear Localization in the Absence of Cell-Cell Contact Is Mediated by a Filamentous Actin-dependent, Myosin II-and Phospho-YAP-independent Pathway during Extracellular Matrix Mechanosensing. J Biol Chem, 2016. 291(12): p. 6096–110.

27. Miralles, F., et al., Actin dynamics control SRF activity by regulation of its coactivator MAL. Cell, 2003. 113(3): p. 329–42.

28. Mouilleron, S., et al., Structure of a pentavalent G-actin*MRTF-A complex reveals how G-actin controls nucleocytoplasmic shuttling of a transcriptional coactivator. Sci Signal, 2011. 4(177): p. ra40.

29. Cui, Y., et al., Yes-associated protein nuclear translocation promotes anabolic activity in human articular chondrocytes. Osteoarthritis Cartilage, 2023. 31(8): p. 1078–1090.

30. Delve, E., et al., CDC42 regulates the expression of superficial zone molecules in part through the actin cytoskeleton and myocardin-related transcription factor-A. J Orthop Res, 2018. 36(9): p. 2421–2430.

31. Gonzalez-Nolde, S., et al., The Actin Cytoskeleton as a Regulator of Proteoglycan 4. Cartilage, 2025. 16(3): p. 376–387.

32. Fath, T., et al., New aspects of tropomyosin-regulated neuritogenesis revealed by the deletion of Tm5NM1 and 2. Eur J Cell Biol, 2010. 89(7): p. 489–98.

33. Stanton, H., et al., ADAMTS5 is the major aggrecanase in mouse cartilage in vivo and in vitro. Nature, 2005. 434(7033): p. 648–52.

34. Schmittgen, T.D. and K.J. Livak, Analyzing real-time PCR data by the comparative C(T) method. Nat Protoc, 2008. 3(6): p. 1101–8.

35. Inguito, K.L., et al., Stress deprivation of tendon explants or Tpm3.1 inhibition in tendon cells reduces F-actin to promote a tendinosis-like phenotype. Mol Biol Cell, 2022. 33(14): p. ar141.

36. West, V.C., et al., Actin Polymerization Status Regulates Tenocyte Homeostasis Through Myocardin-Related Transcription Factor-A. Cytoskeleton (Hoboken), 2025. 82(9): p. 539–555.

37. Patel, S.D., et al., Whole mount staining of lenses for visualization of lens epithelial cell proteins. MethodsX, 2021. 8: p. 101376.

38. Motulsky, H.J. and R.E. Brown, Detecting outliers when fitting data with nonlinear regression - a new method based on robust nonlinear regression and the false discovery rate. BMC Bioinformatics, 2006. 7: p. 123.

39. Masedunskas, A., et al., Parallel assembly of actin and tropomyosin, but not myosin II, during de novo actin filament formation in live mice. J Cell Sci, 2018. 131(6).

40. Bonello, T.T., et al., A small molecule inhibitor of tropomyosin dissociates actin binding from tropomyosin-directed regulation of actin dynamics. Sci Rep, 2016. 6: p. 19816.

41. Liu-Chittenden, Y., et al., Genetic and pharmacological disruption of the TEAD-YAP complex suppresses the oncogenic activity of YAP. Genes Dev, 2012. 26(12): p. 1300–5.

42. Gupta, M., A. Korol, and J.A. West-Mays, Nuclear translocation of myocardin-related transcription factor-A during transforming growth factor beta-induced epithelial to mesenchymal transition of lens epithelial cells. Mol Vis, 2013. 19: p. 1017–28.

43. Hayashi, K., et al., RPEL proteins are the molecular targets for CCG-1423, an inhibitor of Rho signaling. PLoS One, 2014. 9(2): p. e89016.

44. Parreno, J., et al., Tropomyosin 3.1 Association With Actin Stress Fibers is Required for Lens Epithelial to Mesenchymal Transition. Invest Ophthalmol Vis Sci, 2020. 61(6): p. 2.

45. Kee, A.J., et al., On-target action of anti-tropomyosin drugs regulates glucose metabolism. Sci Rep, 2018. 8(1): p. 4604.

46. Kee, A.J., et al., An actin filament population defined by the tropomyosin Tpm3.1 regulates glucose uptake. Traffic, 2015. 16(7): p. 691–711.

47. Shin, H., D. Kim, and D.M. Helfman, Tropomyosin isoform Tpm2.1 regulates collective and amoeboid cell migration and cell aggregation in breast epithelial cells. Oncotarget, 2017. 8(56): p. 95192–95205.

48. Delve, E., et al., YAP/TAZ regulates the expression of proteoglycan 4 and tenascin C in superficial-zone chondrocytes. Eur Cell Mater, 2020. 39: p. 48–64.

49. Campbell, J.J., et al., Loading alters actin dynamics and up-regulates cofilin gene expression in chondrocytes. Biochem Biophys Res Commun, 2007. 361(2): p. 329–34.

50. Jansen, S. and B.L. Goode, Tropomyosin isoforms differentially tune actin filament length and disassembly. Mol Biol Cell, 2019. 30(5): p. 671–679.

