## Supplemental Information for "Tropomyosin 3.1 Regulates Chondrocyte Cortical F-actin Stability and Homeostasis"

**Supplementary Table 1.**
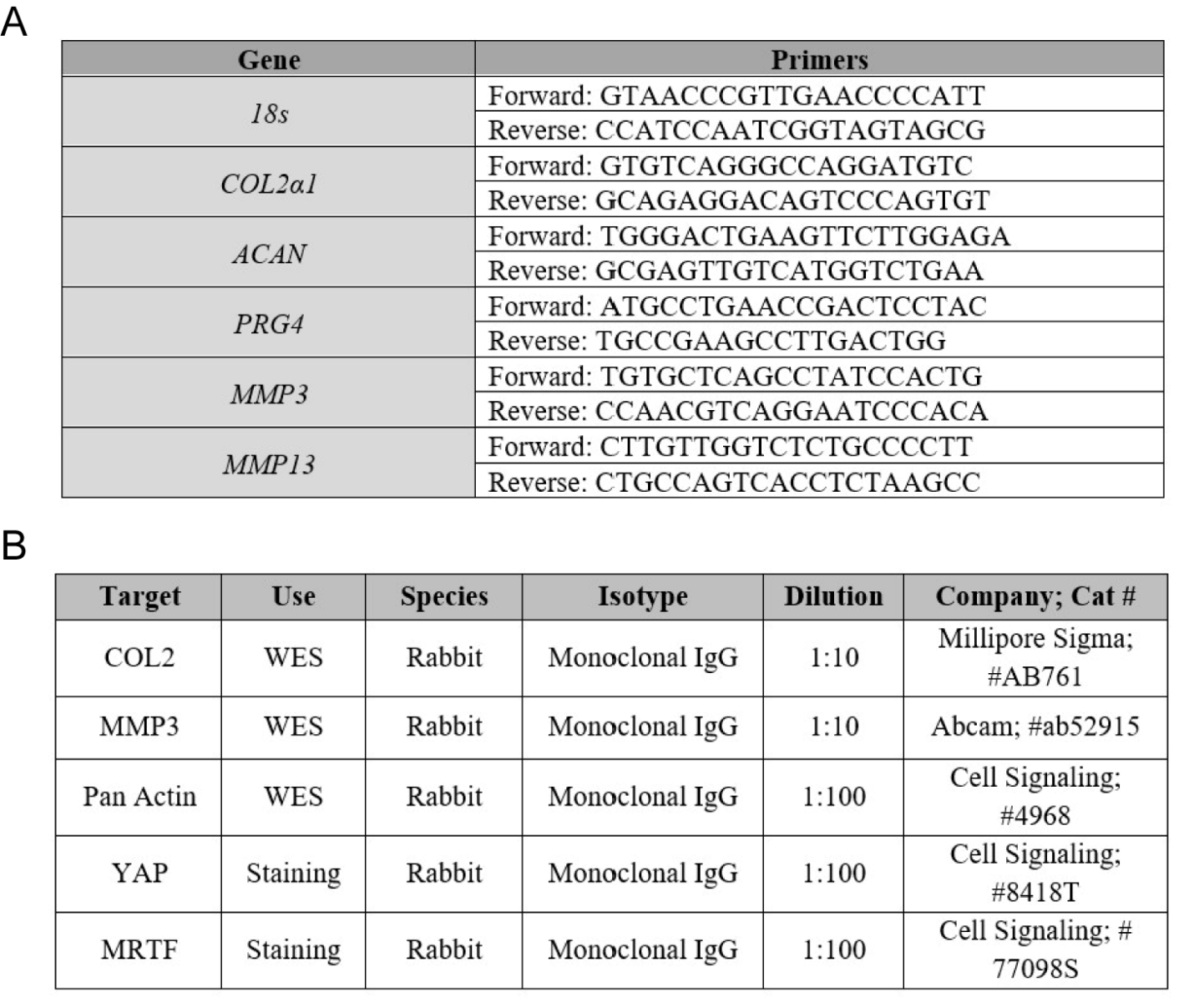

**Supplementary Table 1.** (A) Bovine PCR primers with forward and reverse sequences (B) Primary antibodies used for WES and staining procedures (Abcam; Waltham, MA) (Cell Signaling; Danvers, MA).

**Supplementary Figure 1.**
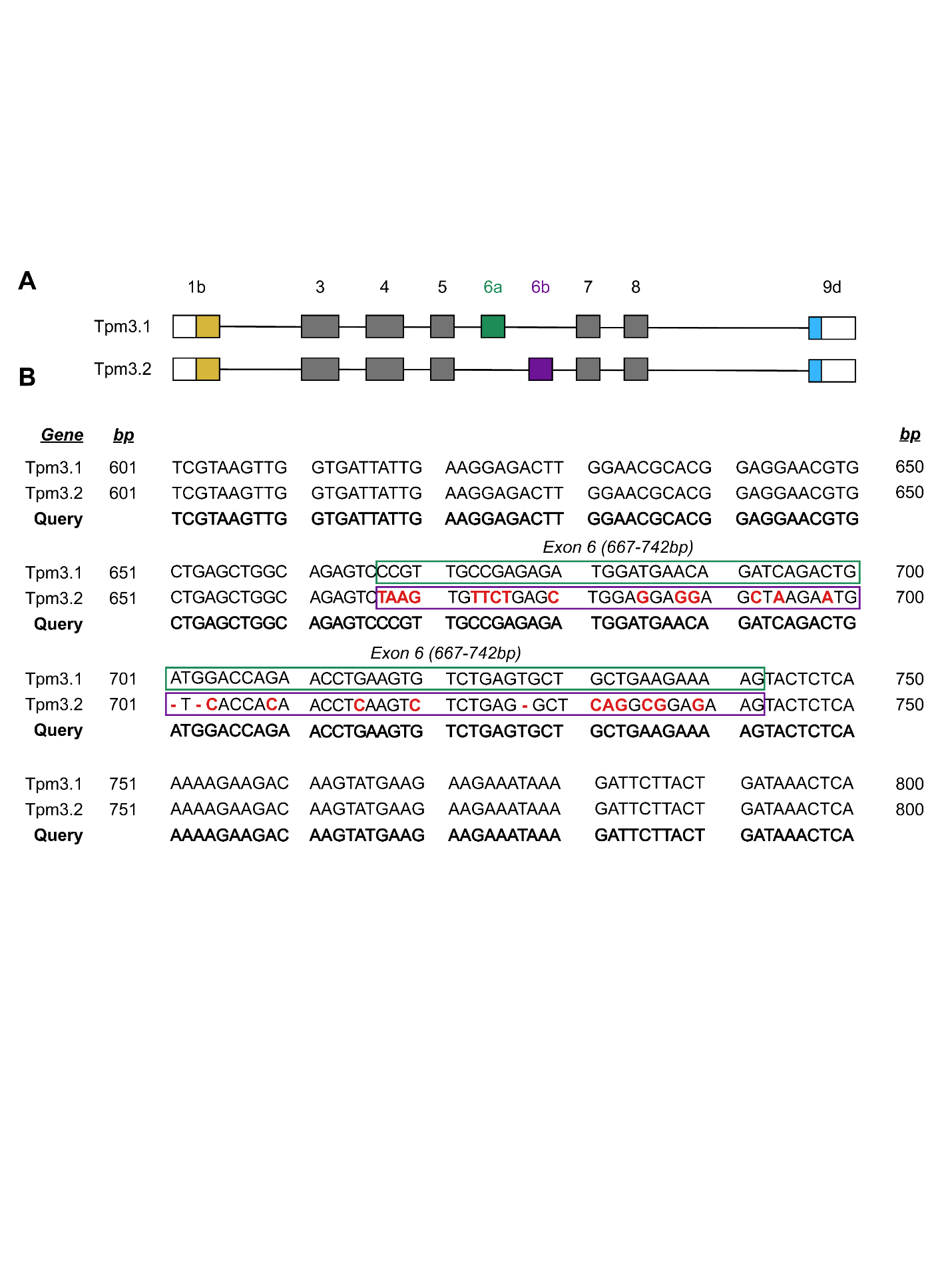

**Supplementary Figure 1. DNA Sequencing Elucidating Differences in Tpm3.1 and Tpm3.2.** (A) Exons for *Tpm3.1* and *Tpm3.2* demonstrating deviations with the presence of 6a (green) vs 6b (purple) exons. (B) Sanger sequencing confirmed *Tpm3.1* based on alignment (green box) with query sequence. Misalignment between query and *Tpm3.2* (purple box) is shown as red-labeled nucleotides, whereas red dashes represent potential nucleotide deletions.

**Supplementary Figure 2.**
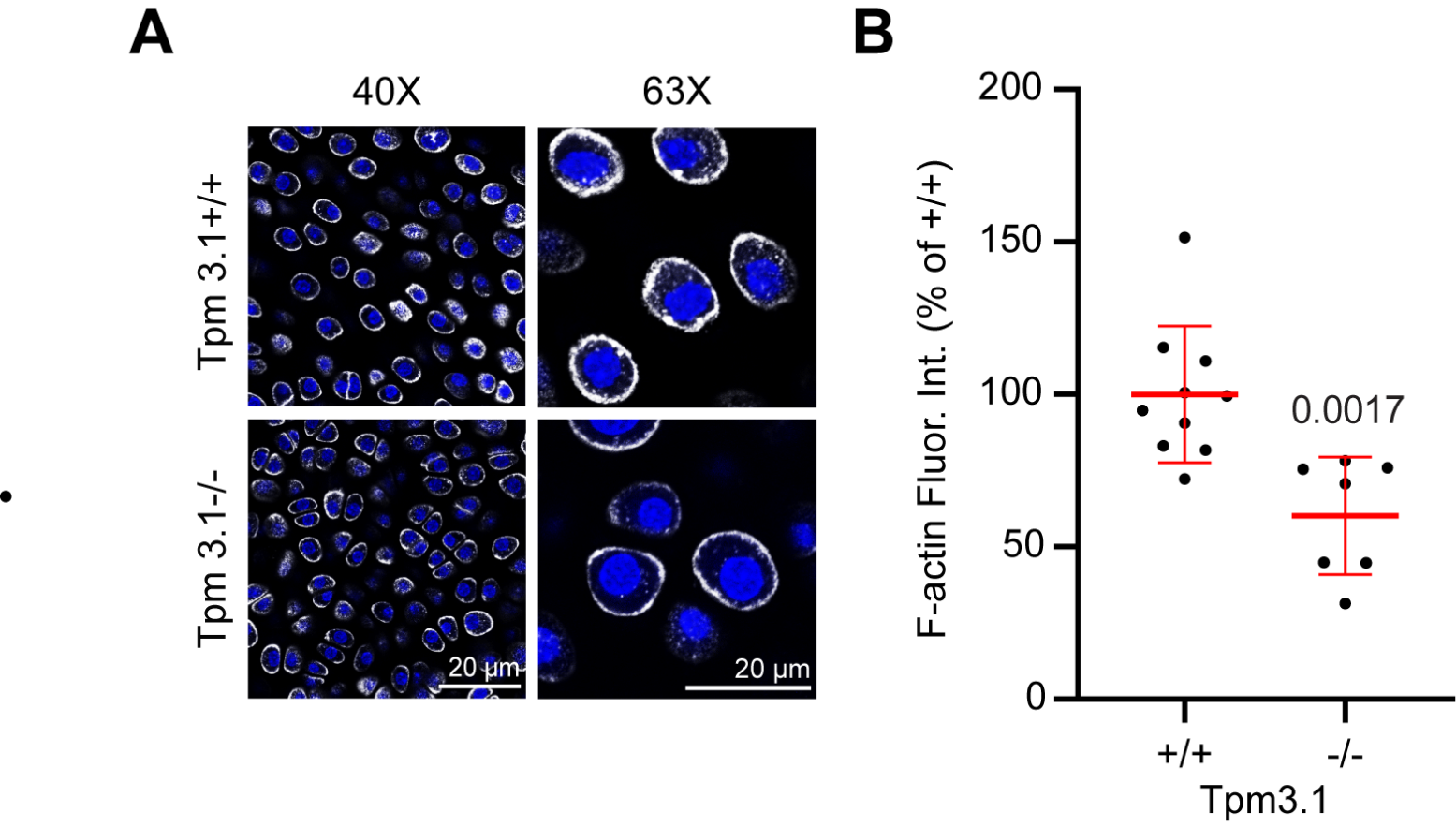

**Supplementary Figure 2. Tpm3.1 Knockout Reduces F-actin.** (A) Confocal microscopy showing F-actin organization and intensity (Grey = phalloidin for F-actin; Blue = Hoechst for nuclei) of Native Tpm3.1 wild-type (Tpm3.1+/+) mouse femoral head chondrocytes (top) and Native Tpm3.1 knockout (Tpm3.1-/-) mouse femoral head chondrocytes (bottom). (B) Dot plot of whole-mount F-actin fluorescence intensity as a percent of wild-type control. Each point represents the average intensity of chondrocytes from one femoral head consisting of between 20-30 chondrocytes. (n= 10 Tpm3.1 +/+ animals, n= 7 Tpm3.1 -/- animals).

**Supplementary Figure 3.**
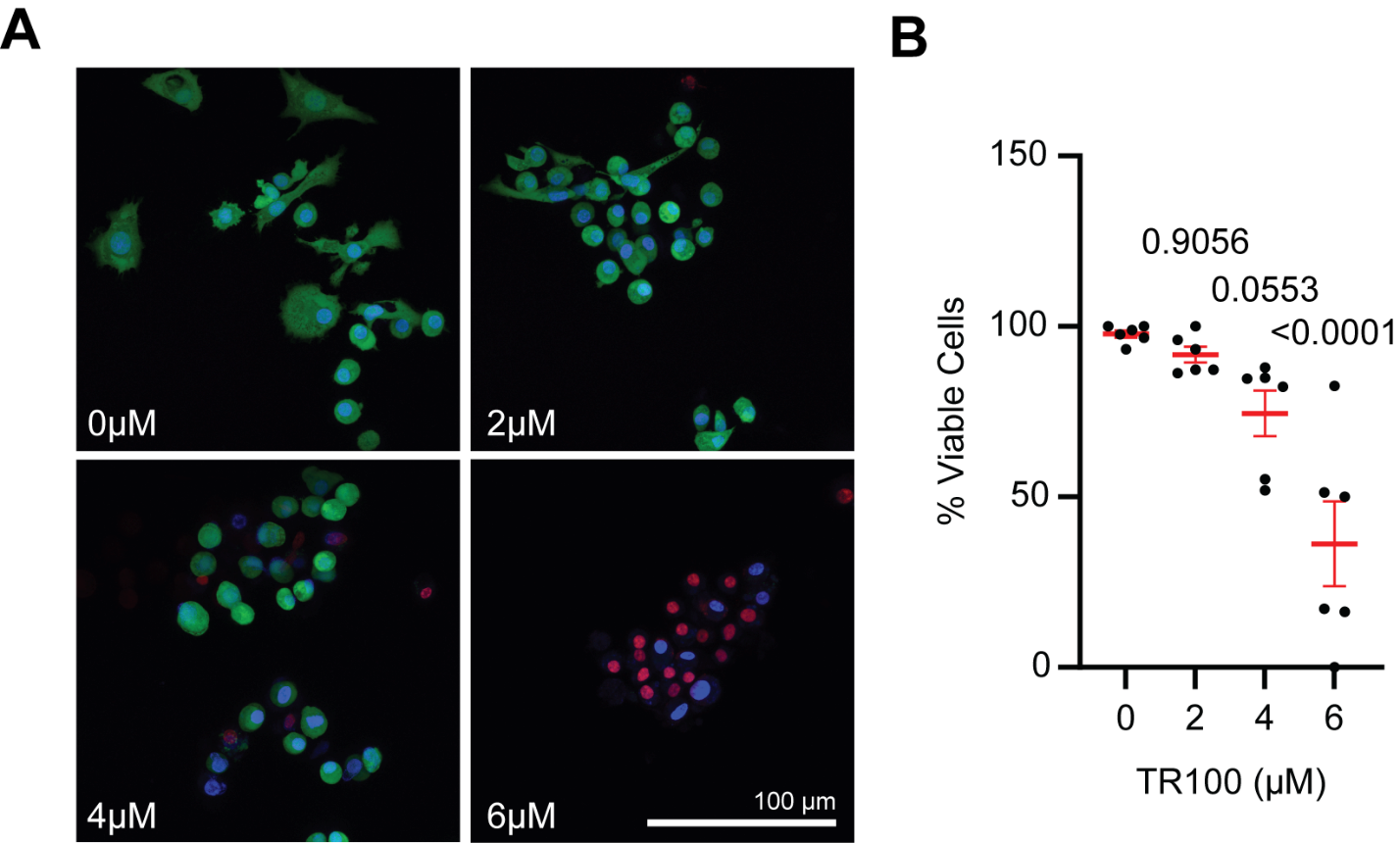

**Supplementary Figure 3. TR100 Does Not Cause Overt Chondrocyte Death.** Chondrocytes treated with TR100 for 24 hours. (A) Confocal images taken at 40X, calcine AM (green) representing live chondrocytes and Ethidium Homodimer (red) representing dead chondrocytes counterstained with Hoechst for nuclei (blue). (B) Quantification of viable cells as a percent of living cells (green) out of total cells (blue). Each point represents an average of 15-20 cells per trial across 6 trials.

**Supplementary Figure 4.**
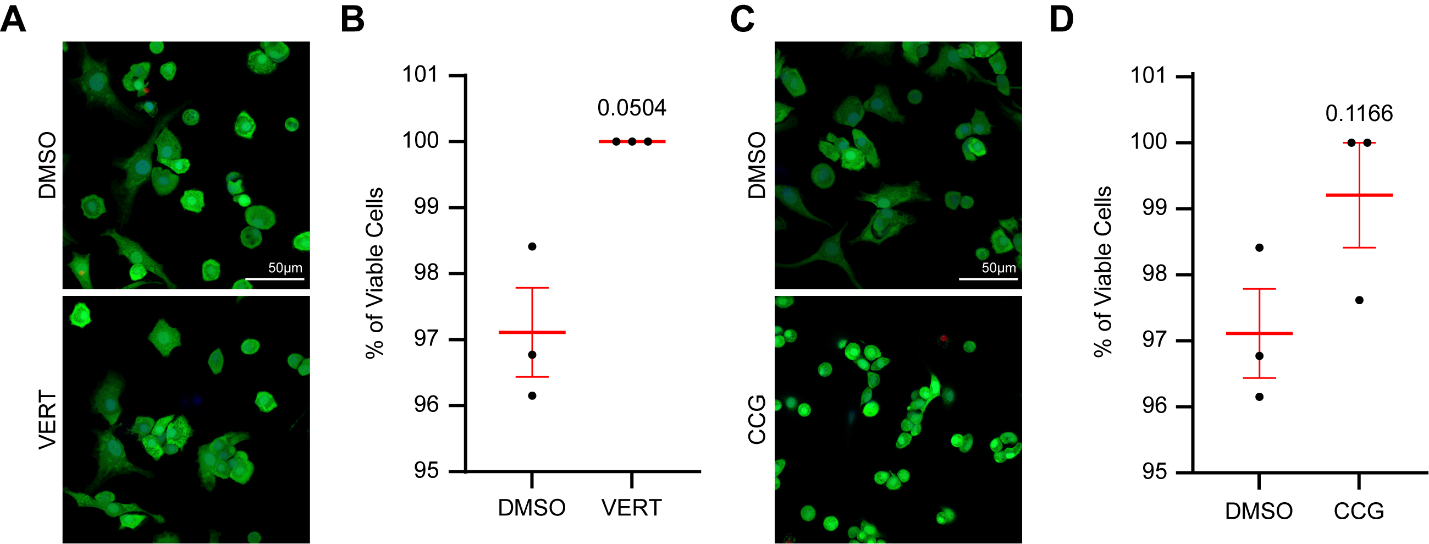

**Supplementary Figure 4. Verteporfin and CCG1423 Do Not Cause Overt Chondrocyte Death.** Chondrocytes treated for 24 hours with (A) 5μM of Verteporfin (VERT) or (C) 10μM of CCG1423 (CCG), confocal images taken at 40X, calcine AM (green) representing live chondrocytes and Ethidium Homodimer (red) representing dead chondrocytes counterstained with Hoechst for nuclei (blue). Quantification of viable cells as a percent of living cells (green) out of total cells (blue) for (B) VERT and (D) CCG. Each point represents an average of 15-20 cells per trial across 3 trials.
